# The Role of the Nucleus Reuniens in Regulating Contextual Conditioning with the Predator Odor TMT

**DOI:** 10.1101/2021.06.10.447925

**Authors:** Laura C. Ornelas, Kalynn Van Voorhies, Joyce Besheer

**Author notes:** Correspondence: Joyce Besheer, Ph.D., Professor, Bowles Center for Alcohol Studies, Department of Psychiatry, University of North Carolina at Chapel Hill, Chapel Hill, NC 27599-7171.

## Abstract

**Rationale:** Experiencing intrusive distressing memories of a traumatic event(s) is a prominent symptom profile for post-traumatic stress disorder (PTSD). Understanding the neurobiological mechanisms associated with this symptom profile can be invaluable for effective treatment for PTSD.

**Objectives:** Here, we investigated the functional role of the nucleus reuniens (RE), a midline thalamic in modulating stressor-related memory.

**Methods:** Female Long Evans rats were implanted with a cannula aimed at the RE. The RE was pharmacologically inactivated via muscimol (0.5 mM) prior to exposure to the predator odor stressor trimethylthiazoline (TMT; synthetically derived fox feces component) or water (controls) in a distinct context with bedding material (Experiment 1) or no bedding (Experiment 2). To measure context reactivity, the index of the contextual memory, 2 weeks following exposure to TMT, rats were re-exposed to the TMT-paired context (in the absence of TMT).

**Results:** In Experiment 1, during context re-exposure (with bedding), inactivation of the RE had no effect on context reactivity. In Experiment 2, during context re-exposure (no bedding), rats previous exposed to TMT showed decreased immobility compared to controls, indicating reactivity to the context and likely related to increased exploration of the environment. Rats in the TMT group that received RE inactivation showed increased immobility relative to rats that received aCSF, suggesting that muscimol pre-treatment blunted context reactivity.

**Conclusion:** In conclusion, recruitment of the RE in stressor-related contextual memory appears to be dependent on the contextual environment and whether the animal is able to engage in different stress coping strategies.

## Introduction

It is estimated that up to 90% of the general population will experience a severe traumatic event during their lifetime (Kessler, Sonnega, Bromet, Hughes, & Nelson, 1995; Yehuda, 2004), which for some could lead to the development of post-traumatic stress disorder (PTSD) (Breslau et al., 1998). Lifetime prevalence of PTSD in the United States is approximately 8.3% for adults, with females (12.8%) being more likely to have a lifetime prevalence of PTSD compared to males (5.7%) (Kessler et al., 2005; Kilpatrick et al., 2013). Further, re-experiencing trauma or memories associated with the traumatic event can lead to increased severity of PTSD symptoms and debilitating long-term effects (APA, 2013).Specifically, one of the most well-known DSM-5 diagnostic criteria for PTSD includes experiencing intrusive distressing memories of the traumatic event(s) (APA, 2013). Understanding the behavioral and neurobiological mechanisms associated with these persistent and long-lasting symptoms can be invaluable tools for effective treatment and prevention strategies for PTSD.

The nucleus reuniens (RE), a midline thalamic nucleus, is an emerging area of the brain underlying symptom profiles of PTSD, such as stress and depression (Kafetzopoulos et al., 2018). The RE receives direct projections from the medial prefrontal cortex (mPFC) (Ramanathan, Jin, Giustino, Payne, & Maren, 2018), while also projecting to the hippocampus; therefore, acting as an interconnected nucleus between these two regions (Eichenbaum, 2017; Jayachandran et al., 2019). As such, the RE has been show to participate in various aspects of fear learning and memory including encoding and retrieval of contextual fear memories after Pavlovian fear conditioning in rats (Ramanathan, Ressler, Jin, & Maren, 2018), as well as extinction of contextual fear memories (Ramanathan, Jin, et al., 2018; Ramanathan & Maren, 2019; Silva, Burns, & Graff, 2019). Therefore, the RE could be an important region to target when investigating neurobiological mechanisms that regulate contextual memory associated with traumatic experiences. In the current study, we utilize an animal model of traumatic stress, inescapable exposure to a predator odor, to investigate the functional role of the RE in modulating stress-related contextual conditioning.

Animal models have become increasingly important in stress research to examine behaviors that can inform our understanding of clinical PTSD symptoms. For example, re-exposing animals to stress-related stimuli or the environment in which the stressor was presented can induce contextual fear or stress responses that can serve as an index of memory of that context. Specifically, predator odor exposure such as bobcat urine produces contextual avoidance to an odor-paired chamber in rats (Albrechet-Souza & Gilpin, 2019; Weera, Schreiber, Avegno, & Gilpin, 2020), and exposure to the synthetically produced predator odor 2,5-dihydro-2,4,5-trimethylthiazoline (TMT; an extract of fox feces), produces contextual stress responses, as evidenced by increased freezing behavior during re-exposure to exposure context (Schwendt et al., 2018; Tyler, Weinberg, Lovelock, Ornelas, & Besheer, 2020). The current study uses TMT exposure as the stressor because it activates a hardwired “learned-independent system” shown to induce innate fear and defensive behaviors (Rosen, Asok, & Chakraborty, 2015). Further, we have previously shown that an important advantage of using predator odor exposure, including TMT, as a stressor, is the ability to measure stress-reactive behaviors during stressor exposure, including defensive digging and immobility (Ornelas, Tyler, Irukulapati, Paladugu, & Besheer, 2020; Tyler et al., 2020). Defensive digging is interpreted as a proactive response to stress (Arakawa, 2007; De Boer & Koolhaas, 2003; Fucich & Morilak, 2018), such that it reflects an active coping response (Ornelas et al., 2020) or fear-related behavior (Riittinen et al., 1986) and predator-stress responsiveness (Neal, Kent, Bardi, & Lambert, 2018). Rats also engage in immobility behavior during TMT exposure, indicative of passive coping behavior and adaptive response to stress (Ornelas et al., 2020; Tyler et al., 2020). Importantly, during re-exposure to the previously paired-TMT context, these behaviors can be quantified as an index of stress-induced contextual memory. As such, this model is an important tool to examine neurobiological mechanisms associated with stress and fear memory induced by stressor exposure.

The current study investigated the functional involvement of the RE in stress-induced contextual memory. In experiment 1, female Long Evans rats were exposed to TMT in a distinct context, with bedding, to allow for engagement in digging, immobility, and avoidance behaviors. In Experiment 2, female rats were exposed to TMT in a distinct context without bedding, only allowing rats to engage in immobility and avoidance behavior. In both experiments, contextual-induced stress reactive behaviors 2 weeks after TMT exposure were measured. Given that previous work has determined a role for the RE in the acquisition of contextual freezing (Ramanathan, Ressler, et al., 2018), we hypothesized that pharmacological inactivation of the RE prior to stressor exposure would impair memory acquisition of the stressor exposure contextual environment, blunting context reactivity behaviors during re-exposure to the stressor-paired context.

## Materials and Methods

### Subjects

Female young adult (arrived at 7-8 weeks old) Long Evans rats (n=85) were used for these experiments. Animals were doubled housed in ventilated cages (Tecniplast, West Chester, PA) upon arrival to the vivarium with ad libitum food and water. Rats were maintained in a temperature and humidity-controlled colony with a 12-hour light/dark cycle (lights on at 07:00). All experiments were conducted during the light cycle. Animals were handled for five days prior to the start of experiments. Animals were under continuous care and monitoring by veterinary staff from the Division of Comparative Medicine at UNC-Chapel Hill. All procedures were conducted in accordance with the NIH Guide for the Care and Use of Laboratory Animals and institutional guidelines.

### Drugs

Muscimol (R&D systems, Minneapolis, Minnesota) was dissolved in saline (0.9%) to produce 0.5 mM muscimol. Doses were chosen based on previous work (Jaramillo, Randall, Frisbee, & Besheer, 2016; Randall et al., 2021) and our own pilot studies.

### Cannulae Implantation Surgery and Verification

One week before behavioral testing, rats were anesthetized with isoflurane (5% for induction, ~2% for maintenance) and placed in a stereotaxic instrument (Kopf Instruments). Rats received unilateral implantation of a 26-gauage guide cannula (Plastics One, Roanoke, VA) at a 10° angle on the midline aimed to terminate 2 mm above the nucleus reuniens of the thalamus (RE; AP −2.0, ML +1.3, DV −5.8). The guide cannula was affixed to the skull with dental cement, and a dummy injector that did not extend past the guide was secured with a dust cover to prevent debris from entering the cannula. Rats were singled housed after surgery and underwent a 7 d recovery period prior to behavioral testing. Rats received 3 doses of the analgesic Banamine (2.5 mg/kg; 12-16 hr prior to surgery, during recovery and 24 hr later).

At the end of the experiments (see Figure 1 for timeline of experiments), rats were deeply anesthetized with pentobarbital and perfused with 0.1 M PBS, followed by 4% paraformaldehyde, 4°C; pH=7.4. Brains were extracted and stored in 4% paraformaldehyde for 24 hr then transferred to 30% (w/v) sucrose in 0.1M PBS solution. Brains were sliced on a freezing microtome into 40 μm coronal sections and tissue slices were stored in cryoprotectant (Jaramillo et al., 2016). Brain tissue was stained with cresyl violet to verify cannula placement (Figure 2a, b). Only data from rats with cannula/injector tract determined to be in the nucleus reuniens were used in the analyses. Cannula placements we were unable to visually confirm (Experiment 1, n=10/40; Experiment 2, n=4/45) were considered misses and were not included in any data analyses.

**Figure 1.**
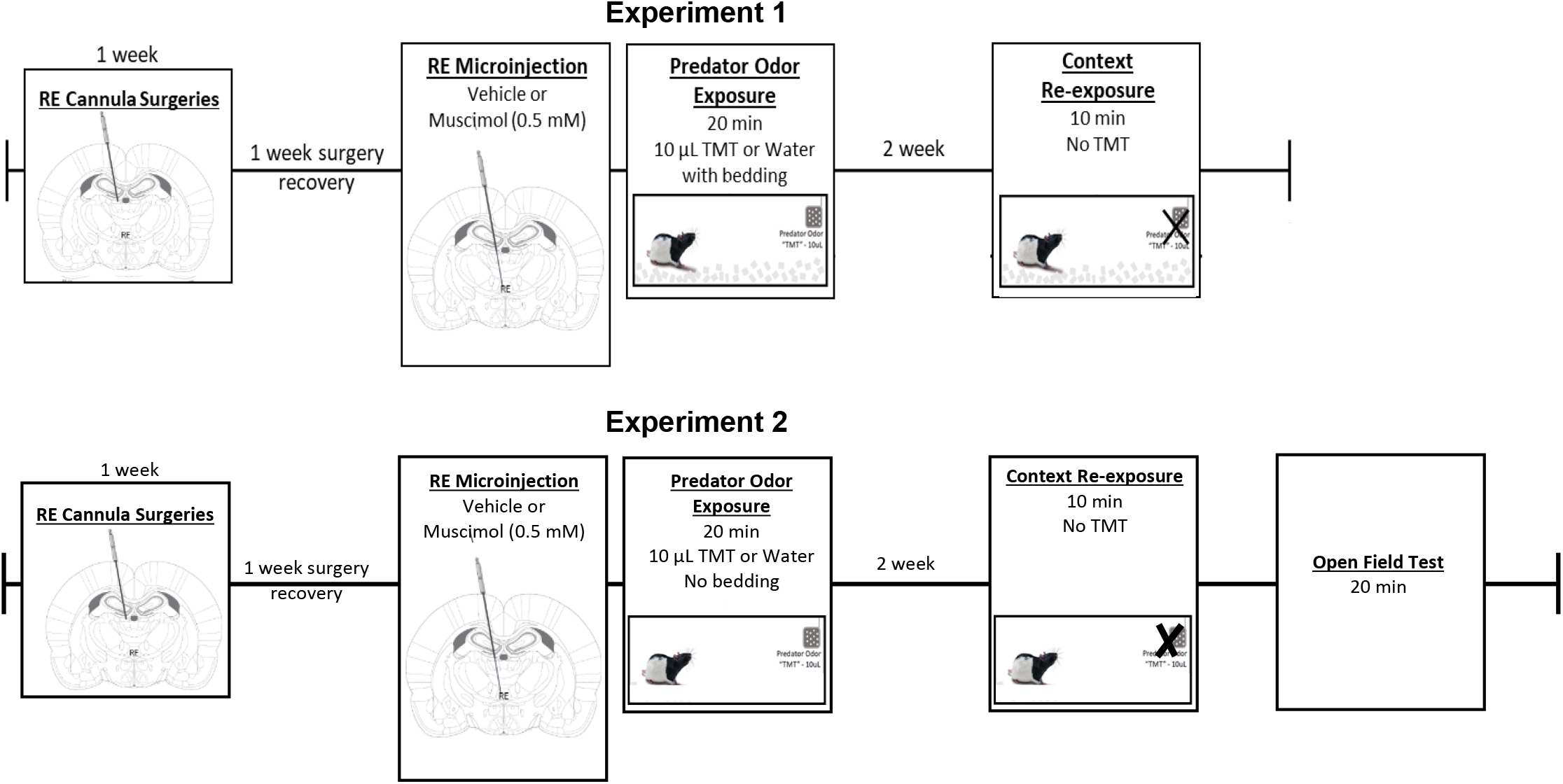
Timeline of Experiments. Experiment 1: Female Long-Evans rats received unilateral cannula (2 mm dorsal RE). After 1 week of surgery recovery, rats received microinjections of vehicle or muscimol (0.5mM) prior to water/TMT exposure (with bedding). 2 weeks later, rats were re-exposed to the initial TMT-paired contextual environment in the absence of TMT. The sample size of each treatment group is as followed: aCSF, water (n=10); muscimol, water (n=8); aCSF, TMT (n=8); muscimol, TMT (n=8). Experiment 2: *Surgery and microinjection procedures same as Experiment.* Water/TMT exposure occur without bedding for Experiment 2. Two weeks later, rats were re-exposed to the initial TMT-paired contextual environment in the absence of TMT. The sample size of each treatment group is as followed: aCSF, water (n=5); muscimol, water (n=7); aCSF, TMT (n=7); muscimol, TMT (n=9).

**Figure 2a.**
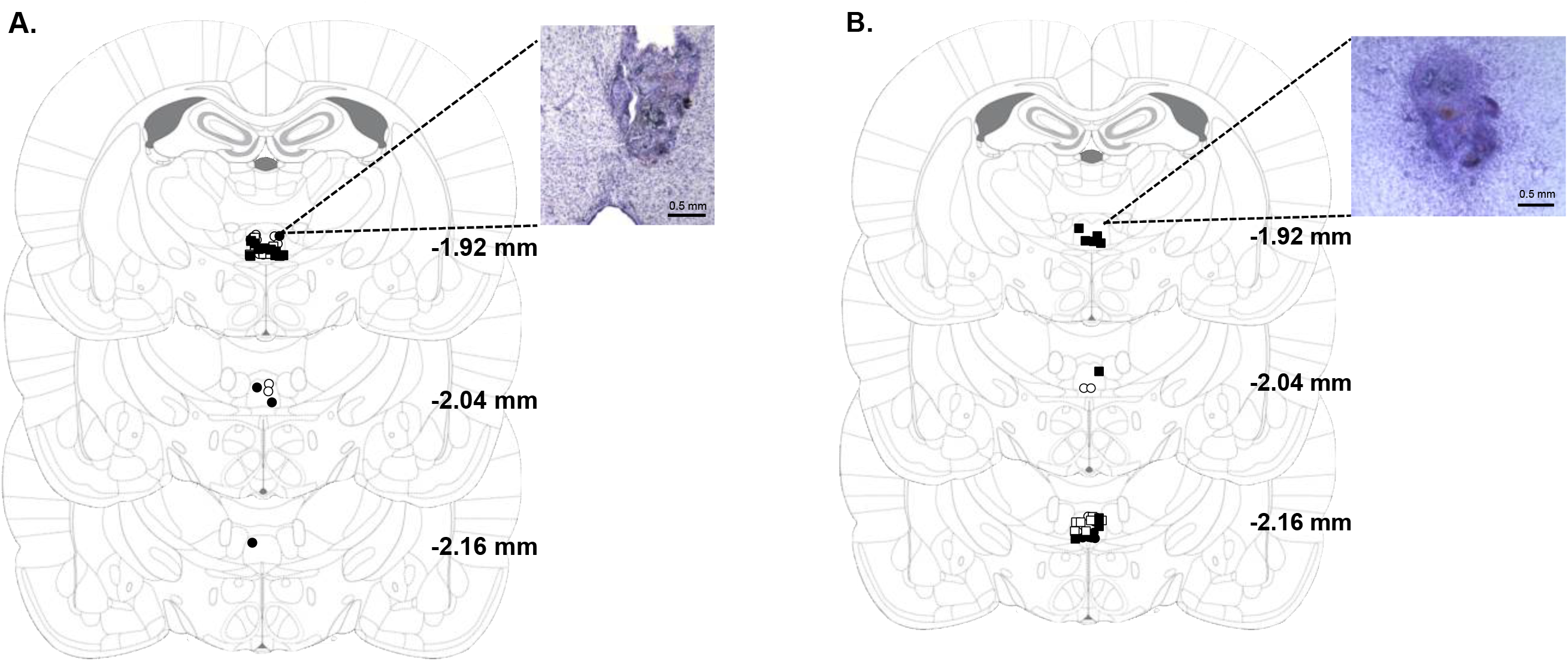
Experiment 1: Unilateral RE cannula placements and representative photomicrograph (4X). Symbols representing RE hits for each treatment group are as followed: open circles, aCSF, water; closed circles, muscimol, water; open squares, aCSF, TMT; closed squares, muscimol, TMT. **Figure 2b.** Experiment 2: Unilateral RE cannula placements and representative photomicrograph (4X). Symbols representing RE hits for each treatment group are as followed: open circles, aCSF, water; closed circles, muscimol, water; open squares, aCSF, TMT; closed squares, muscimol, TMT.

### Microinjection Procedures

For intracranial microinjections prior to predator odor exposure, rats were transferred in the home cage to a separate room from the TMT exposure room. A stainless steel injector (33 gauge) connected to polyethylene tubing was inserted into the guide cannula for drug infusions. Tubing was connected to a Hamilton syringe (5 μL), which was connected to an infusion pump. For both experiments 1 and 2, vehicle (aCSF) and muscimol (0.5 mM) microinjections were delivered through an injector extending 2 mm below the guide cannula at a volume of 0.3 μL across 1 min. The injector remained in place for an additional 2 min after the infusion to allow for diffusion. Water or TMT exposure sessions occurred approximately 7-10 min after the end of the diffusion period.

### Predator Odor Exposure and Context Re-exposure

Following aCSF or muscimol infusion, rats were transported to the testing room. Rats were transferred from the home cage to Plexiglas exposure chambers (45.72 ×17.78 ×21.59 cm; UNC Instrument Shop, Chapel Hill, NC, USA). The length of the back wall of the chambers was opaque white with two opaque black side walls and a clear front wall to allow for video recordings. A metal basket (17.8 cm above the floor) piece of filter paper on which was placed 10 μl of TMT or water (for controls) so that the filter paper was inaccessible to the rat. Note: all water exposures occurred before the TMT exposures to ensure there was not TMT odor in the room during water exposure sessions. For experiment 1, approximately 600 mL of white bedding (Shepherds ALPHA-dri) was added to the bottom of the exposure chamber prior to the animal being placed in the chamber. For Experiment 2, no bedding was added to bottom of the chamber. After the rat was placed into the chamber, a clear Plexiglas top was slid and secured into place. The exposure session was 20 min in duration and recorded by a video camera for later analysis using ANY-maze ™ Video Tracking System (Stoelting Co. Wood Dale, IL, USA). Following the exposure session, each rat was returned to the homecage.

#### Context re-exposure

Two weeks following predator odor exposure, animals were placed in the context in which they had been previously exposed to water or TMT for 10 min (no TMT present) and behavior was monitored.

### Locomotor Activity

To confirm that the muscimol dose produced no effects on locomotor behavior, a subset of rats from Experiment 2 was injected with vehicle and muscimol (0.5 mM) into the RE three weeks after the initial TMT Exposure. Approximately 7-10 min after infusion, rats were placed in an open field for monitoring of general locomotor activity for 20 min. Open field chambers (44.4 × 22.9 × 30.5 cm; Med Associates Inc.; St. Albans, VT) were individually located within sound-attenuating cubicles equipped with an exhaust fan that provided both ventilation and masking of external sounds. Time and distance spent on each side of the chamber was measured with 4 parallel beams across the chamber floor.

### Data Analyses

#### TMT and Context Re-exposure

Using ANY-maze software, the length of the rectangular TMT exposure chamber was divided into two sides for analysis (TMT side and non-TMT side) to allow for analysis of time spent on the side in which the TMT was located. Digging behavior was quantified manually from the video recording. Immobility was operationally defined as lack of movement for more than 2 seconds as assessed using ANY-maze software. Therefore, immobility likely captures both inactivity and freezing, which is characteristic of a fear response in rodents and observed during TMT exposure (Ornelas et al., 2020). Two-way ANOVA was used to compare group differences in stress-reactive behaviors (digging, immobility, time spent on TMT side) during TMT exposure and context re-exposure. Time spent digging, immobile and time spent on TMT side were analyzed by three-way RM ANOVA with TMT exposure and drug treatment as between-subjects factor and time as a within-subjects factor. Tukey multiple comparisons tests were used to follow up significant main effects of groups and interactions. All data are reported as mean ± S.E.M. Significance was set at *p* ≤ 0.05.

#### Locomotor Activity

T-tests were used to compare differences in locomotor activity (distance traveled) in vehicle and muscimol treated animals. All data are reported as mean ± S.E.M. Significance was set at *p* ≤ 0.05.

## Results

### Experiment 1: Pharmacological Inactivation of Nucleus Reuniens Does Not Affect Active or Passive Coping Behaviors during TMT Exposure or Context Re-exposure

During TMT exposure (with bedding), rats engaged significantly greater defensive digging (Figure 3A, F (1,30) = 19.77, *p* < 0.05), immobility behavior (Figure 3B, F (1,30) = 46.07, *p* < 0.05), and avoidance of the TMT side (Figure 3C, F(1,30) = 16.59, *p* < 0.05) compared to controls. Examination of fecal boli showed a significant main effect of TMT (F(1,30) = 16.55, *p* < 0.05) with more fecal boli compared to controls (Veh, Water: 0.70 ± 0.42; Muscimol, Water: 1.00 ± 0.33; Veh, TMT: 4.00 ± 1.02; Muscimol, TMT: 3.00 ± 0.87). There was no significant main effect of drug (*p* > 0.05), indicating that pharmacological inactivation of nucleus reuniens did not significantly alter active or passive coping behaviors during TMT exposure. There was no significant drug x exposure interaction (*p* > 0.05).

**Figure 3.**
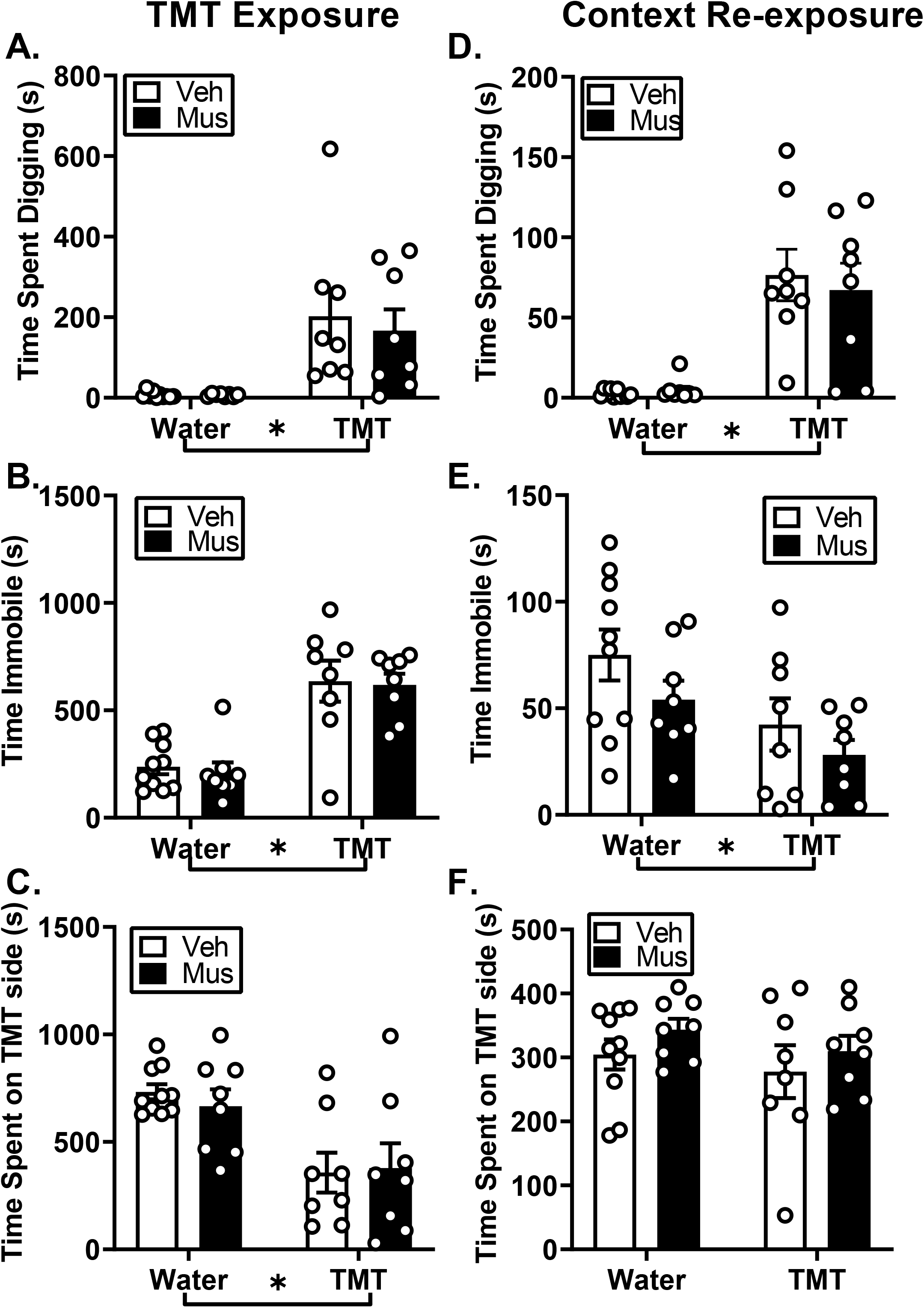
Effects of TMT exposure on active and passive coping behaviors and context reactivity in female rats. All female rats exposed to TMT (with bedding) engaged in defensive digging (**A**), immobility (**B**) and avoidance (**C**), but RE inactivation did not alter active or passive coping behaviors during TMT exposure. During context re-exposure with bedding, inactivation of the RE had no effect on context re-exposure behavior, but rats showed reactivity to TMT-paired context through increased defensive digging (**D**) and immobility (**E**) but not avoidance, (**F**). Mean ± SEM. * by group labels = main effect of group

During context re-exposure, prior inactivation of the RE had no effect on context re-exposure behavior. That is, rats previously exposed to TMT showed increased defensive digging and decreased immobility behavior (Digging: Figure 3D, F(1,30) = 38.39, *p* < 0.05; Immobility: Figure 3E, F(1,30) = 7.59, *p* < 0.05), indicative of context reactivity, but there were no differences in avoidance behavior and no main effect of drug treatment or significant interaction (Figure 3F, *p* < 0.05). Examination of fecal boli showed no significant main effects of drug, exposure or drug x exposure interaction (p > 0.05) (Veh, Water: 4.5 ± 2.86; Muscimol, Water: 1.25 ± 0.62; Veh, TMT: 0.87 ± 0.62; Muscimol, TMT: 0.75 ± 0.62).

### Experiment 2: Pharmacological Inactivation of Nucleus Reuniens Does Not Affect Passive Coping Behaviors during Predator Odor Exposure but Blunts Context Reactivity

During TMT exposure, rats engaged in significantly greater stress-reactive behaviors including immobility behavior (Figure 4A, F(1,24) = 139.5, *p* < 0.05) avoidance (Figure 4B, F(1,24) = 66.43, *p* < 0.05) compared to controls. There was a significant main effect of drug treatment (Immobility: Figure 4A, F(1,24) = 6.65; Avoidance: Figure 4B, F(1,24) = 4.33, *p* < 0.05); however, analysis comparing vehicle and muscimol groups showed no significant differences between immobility and avoidance (*p* > 0.05). There was no significant exposure x drug interaction (*p* < 0.05), indicating that pharmacological inactivation of nucleus reuniens did not significantly alter stress-reactive behaviors during TMT exposure. Examination of fecal boli showed a significant main effect of TMT (F(1,24) = 4.11, *p* ≤ 0.05) showing more fecal boli compared to controls (Veh, Water: 1.33 ± 0.67; Muscimol, Water: 1.57 ± 0.65; Veh, TMT: 4.00 ± 1.46; Muscimol, TMT: 2.63 ± 0.53).

**Figure 4.**
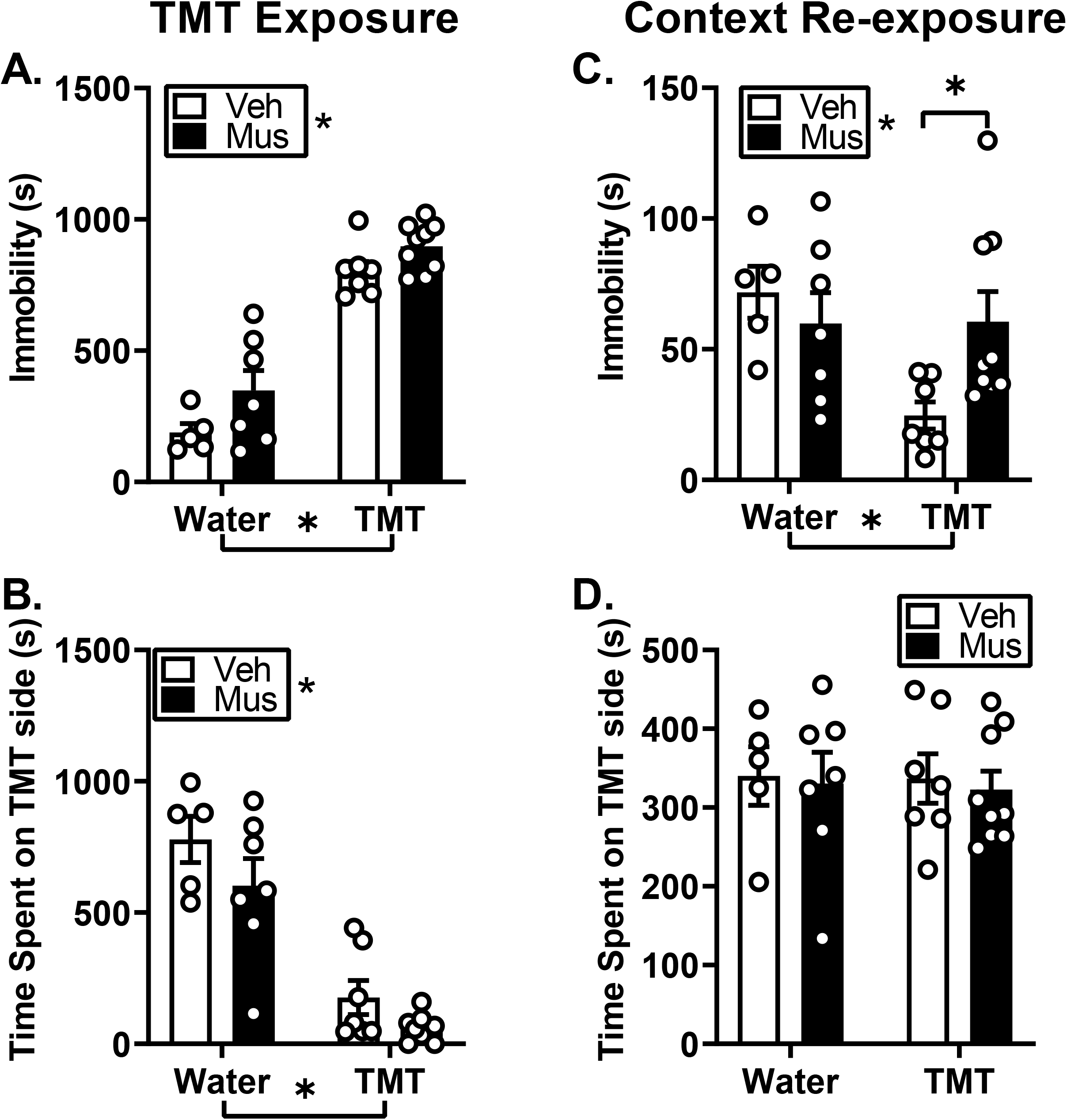
Effects of TMT exposure on passive coping behaviors and context reactivity in female rats. All female rats exposed to TMT (without bedding) engaged in immobility (**A**) and avoidance (**B**), but RE inactivation did not alter passive coping behaviors during TMT exposure. During context re-exposure rats in the TMT group that received RE inactivation showed increased immobility relative to rats that received aCSF (**C**). Rats did not show avoidance behavior and RE inactivation did not alter avoidance behavior (**D**). Mean ± SEM. * by group labels = main effect of group

During context re-exposure, a two-way ANOVA revealed a significant exposure x treatment interaction (Figure 4C, F (1,24) = 4.97, *p* < 0.05) for immobility behavior. Rats in the TMT group that received RE inactivation showed increased immobility relative to rats that received aCSF (4C, *p* < 0.05), suggesting that muscimol pre-treatment blunted context reactivity. Analysis of avoidance behavior during context re-exposure showed no main effect of exposure, drug or significant time x exposure x drug interaction (Figure 4D, *p* > 0.05). Examination of fecal boli showed no significant main effect of exposure, drug or exposure x drug interaction (Veh, Water: 2.00 ± 1.05; Muscimol, Water: 2.00 ± 1.14; Veh, TMT: 1.29 ± 0.64; Muscimol, TMT: 1.44 ± 0.60).

### Experiment 3: Muscimol (0.5 mM) Produced No Effects on Locomotor Behavior

To confirm drug treatments produces no effects on locomotor behavior, we performed microinjection of vehicle or muscimol (0.5 mM) into the RE in a subset of animals. T-tests revealed no main effects of pharmacological treatments (p > 0.05; Figure 5), confirming that this dose of muscimol (0.5 mM) into the RE did not produce an impairment in locomotor behavior.

**Figure 5.**
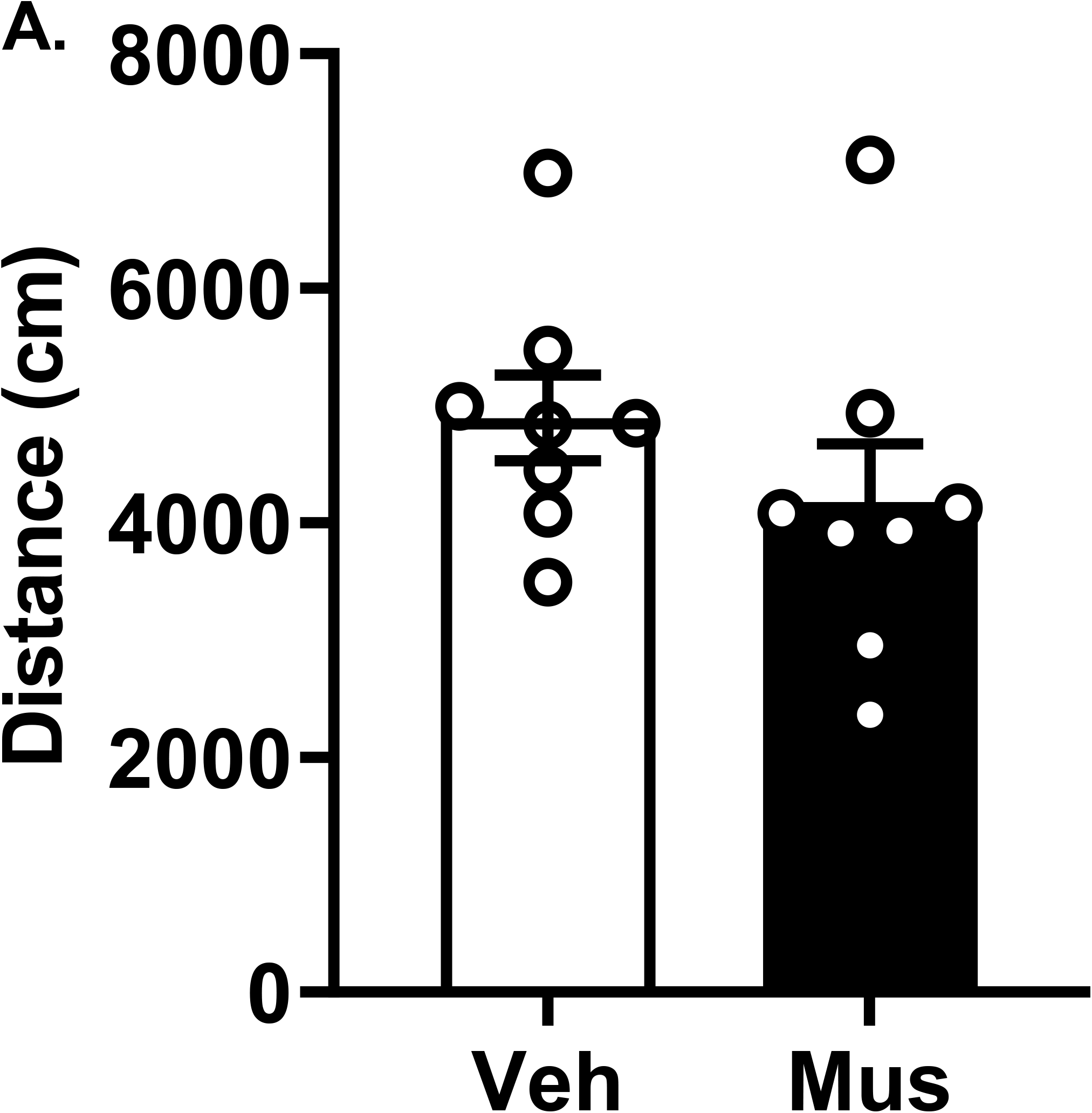
Confirmation drug treatments does not affect locomotion. Open Field Test. Microinjections of vehicle or muscimol (0.5 mM) into the RE produced no differences in locomotor activity between treatments as well as no impairments in overall locomotor activity.

## Discussion

In the current study, we sought to determine the role of the RE in regulating contextual conditioning in response to predator odor stressor, TMT, in female rats. The results support several important findings. First, RE inactivation prior to TMT exposure, does not alter the engagement in stress-reactive behaviors (defensive digging, immobility or avoidance), indicating that the RE is not involved in regulating these behaviors during stressor exposure. Second, RE inactivation blunts context reactivity in rats previously exposed to TMT, and this is dependent on the context of the TMT exposure. This suggests that recruitment of the RE in stressor-related contextual memory is dependent on the contextual environment and may be related to the engagement of different stress coping strategies that are permitted by the environment. Together, these results support a functional role for the RE in stressor-related memory which make it an important region of investigation in the neurobiological mechanisms of stress memory.

The role of the RE has been implicated in fear learning and memory including encoding, acquisition and retrieval of contextual fear memories (Maisson, Gemzik, & Griffin, 2018; Mei, Logothetis, & Eschenko, 2018; Ramanathan, Ressler, et al., 2018). Therefore, we hypothesized RE involvement in the acquisition of a TMT stressor-related memory. First, we show that pharmacological inactivation of the RE does not affect the engagement in stress-reactive behaviors (digging, immobility or avoidance) during the TMT exposure in Experiments 1 and 2. Interestingly, previous research shows that exposure to acute stress (swim and restraint stress) produces high levels of *c-fos* mRNA expression in the RE (Cullinan, Herman, Battaglia, Akil, & Watson, 1995; Kafetzopoulos et al., 2018), suggesting possible recruitment of the region during a stressor. It is known that the RE is recruited during encoding of contextual information during fear conditioning (Lin, Chiou, & Chang, 2020), suggesting this region is involved in the formation and acquisition of stressor or context-related memory (Eichenbaum, 2017). Therefore, we show a potential dissociation between the role of the RE in engagement in stress reactivity behaviors during the TMT predator odor exposure and its role in the memory acquisition of the TMT exposure context.

Re-exposing animals to stress-related stimuli or the environment in which the stressor was presented can induce contextual fear or stress responses that can serve as an index of memory of that context (Ornelas et al., 2020). In the present work, during re-exposure to the previously TMT-paired context, rats engaged in behavioral responses, such as defensive digging and decreased immobility to a greater extent than controls, indicative of context reactivity. This finding is consistent with previous work (Ornelas et al., 2020; Shallcross et al., 2019; Tyler et al., 2020). In Experiment 1, in which bedding was provided in the context, this was evidenced by increased digging and reduced immobility behavior (i.e., greater mobility) relative to controls during the context re-exposure. In Experiment 2, in which no bedding was provided in the context, rats also showed reduced immobility behavior (i.e., greater mobility). We have previously hypothesized that this behavioral profile of increased mobility could represent hypervigilance upon return to the previously paired TMT context (Tyler et al., 2020). Furthermore, avoidance of the TMT-paired side of the chamber was not observed during context re-exposure in either experiment. This is not altogether surprising given that it has been shown that although TMT produces significant avoidance behavior during exposure, TMT has not been shown to induce context conditioning through avoidance behavior (Ornelas et al., 2020; Rosen et al., 2015). Additionally, these data are in accordance with previous work from our lab (Ornelas et al., 2020) and others (Schwendt et al., 2018) that show TMT can produce contextual conditioning.

Next, we examined the functional involvement of the RE in stressor-related memory via inactivation of the RE prior to stressor exposure. In Experiment 1, in which bedding was present in the context, inactivation of the RE had no effect on behavior during context re-exposure. That is, rats engaged in digging behavior and decreased immobility behavior (e.g., hypervigilance) at levels comparable to controls. This outcome suggests that under these conditions the RE is not involved in regulating contextual conditioning. In contrast, in Experiment 2, in which no bedding was present in the context, RE inactivation blunted context reactivity. That is, rats in the TMT group that received intra-RE muscimol showed increased immobility relative to the TMT group that received aCSF, suggesting a role for the RE in regulating contextual conditioning. Together, these results suggest recruitment of the RE in contextual conditioning appears to be dependent on the contextual environment.

There are a few potential explanations as to why the RE is involved in blunting contextual reactivity, specifically involving immobility behavior but not digging behavior. First, in Experiment 1, by providing bedding during TMT exposure and context re-exposure, rats were provided the opportunity to engage in different stress coping strategies including defensive digging and immobility behavior. Engagement in digging behavior may act as a competing behavioral response, inherently activating systems that regulate digging, and override engagement in immobility behavior. Further, engagement in digging behavior may amplify the salience of the environment such that a return to the environment re-engages those systems and this supersedes RE-related circuitry.

In contrast, in Experiment 2, in which the rats primarily engage in immobility during TMT exposure, because there is no bedding in the environment, inactivation of the RE prior to TMT exposure attenuates contextual conditioning. This outcome may be due to an impairment in the ability of the RE to encode the information of the contextual environment. We theorize that the RE plays an integral role in modulating contextual memory of environments which promote immobility behavior compared to environments which allow animals to engage in a other stress-reactive behaviors (e.g., digging). For example, the RE has been implicated in various components of contextual conditioning (Eichenbaum, 2017). Pharmacological inactivation of the RE has been shown to prevent the acquisition of contextual freezing during Pavlovian fear conditioning (Ramanathan, Ressler, et al., 2018), as well as extinction of contextual fear memories (i.e., increased freezing responses during extinction tests) (Ramanathan & Maren, 2019). Inactivation of the RE also produces overgeneralization of contextual fear memory in mice (i.e., increases in freezing in an altered context similar to conditioning context) (Xu & Sudhof, 2013). In the same study, optogenetic stimulation using phasic and tonic patterns produces differential effects on freezing during altered context, further emphasizing the role of the RE in modulating contextual fear memory (Xu & Sudhof, 2013). Therefore, considering the RE is known to specifically regulate freezing or immobility behavior during fear conditioning studies, the results from the current study further illustrate the role of the RE in regulating contextual conditioning.

Neural circuitry associated with the RE also plays a role in modulating contextual conditioning. During encoding of contextual information during fear conditioning (Lin et al., 2020), the RE coordinates mPFC–hippocampal interactions (Eichenbaum, 2017). Further, connectivity between the medial prefrontal cortex (mPFC) and RE may be distinctly involved in regulating immobility behavior during stressor exposure and re-exposure to the stressor-paired context. For example, the prelimbic (PrL) but not infralimbic (IL) cortex has been implicated in modulating freezing behavior during TMT exposure (Fitzpatrick, Knox, & Liberzon, 2011; Hwa et al., 2019). In addition, the mPFC influences memory retrieval through projections to RE which integrates information with the hippocampus (Eichenbaum, 2017; Furtak, Wei, Agster, & Burwell, 2007; Jayachandran et al., 2019). Therefore, these interactions between brain regions would facilitate encoding of a contextual representation of the TMT-paired context to produce contextual conditioning. However, in the current study, it is possible that by pharmacological inactivation of the RE prior to the TMT exposure specifically in a context which promotes immobility behavior, we are preventing the integration of input and output information of the mPFC-RE- hippocampus pathway, and ultimately preventing contextual conditioning.

In conclusion, the current study shows that RE inactivation blunts contextual conditioning (as measured by immobility behavior) in rats previously exposed to TMT when no bedding is present in the context. This suggests that the RE is important for the expression of this context-induced stress memory. However, when providing rats with bedding during TMT exposure, RE inactivation does not affect the expression of the context-induced memory. Together, these data suggest recruitment of the RE in stressor-related contextual memory appears to be dependent on the contextual environment and whether the animal is able to engage in different stress coping strategies (digging and immobility). Overall, these results add to the current literature implementing a role in the RE as an emerging area of the brain underlying symptom profiles of PTSD, specifically re-experiencing of memories associated with trauma exposure.

## Acknowledgement

This work was supported in part by the National Institute of Health AA026537 and AA011605 (JB) and by the Bowles Center for Alcohol Studies. LCO was supported by Diversity Supplement to AA026537. The authors thank Abigail Garcia-Baza for their help with behavioral analysis.

## References

Albrechet-Souza, L., & Gilpin, N. W. (2019). The predator odor avoidance model of post-traumatic stress disorder in rats. Behav Pharmacol, 30(2 and 3-Spec Issue), 105–114. doi:10.1097/FBP.0000000000000460

APA. (2013). Diagnostic and statistical manual of mental disorders (5th ed.). Washington, DC.

Arakawa, H. (2007). Ontogeny of sex differences in defensive burying behavior in rats: effect of social isolation. Aggress Behav, 33(1), 38–47. doi:10.1002/ab.20165

Breslau, N., Kessler, R. C., Chilcoat, H. D., Schultz, L. R., Davis, G. C., & Andreski, P. (1998). Trauma and posttraumatic stress disorder in the community: the 1996 Detroit Area Survey of Trauma. Arch Gen Psychiatry, 55(7), 626–632. doi:10.1001/archpsyc.55.7.626

Cullinan, W. E., Herman, J. P., Battaglia, D. F., Akil, H., & Watson, S. (1995). Pattern and time course of immediate early gene expression in rat brain following acute stress. Neuroscience, 64(2), 477– 505.

De Boer, S. F., & Koolhaas, J. M. (2003). Defensive burying in rodents: ethology, neurobiology and psychopharmacology. Eur J Pharmacol, 463(1-3), 145–161. doi:10.1016/s0014-2999(03)01278-0

Eichenbaum, H. (2017). Prefrontal-hippocampal interactions in episodic memory. Nat Rev Neurosci, 18(9), 547–558. doi:10.1038/nrn.2017.74

Fitzpatrick, C. J., Knox, D., & Liberzon, I. (2011). Inactivation of the prelimbic cortex enhances freezing induced by trimethylthiazoline, a component of fox feces. Behav Brain Res, 221(1), 320–323. doi:10.1016/j.bbr.2011.03.024

Fucich, E. A., & Morilak, D. A. (2018). Shock-probe Defensive Burying Test to Measure Active versus Passive Coping Style in Response to an Aversive Stimulus in Rats. Bio Protoc, 8(17). doi:10.21769/BioProtoc.2998

Furtak, S. C., Wei, S. M., Agster, K. L., & Burwell, R. D. (2007). Functional neuroanatomy of the parahippocampal region in the rat: the perirhinal and postrhinal cortices. Hippocampus, 17(9), 709–722. doi:10.1002/hipo.20314

Hwa, L. S., Neira, S., Pina, M. M., Pati, D., Calloway, R., & Kash, T. L. (2019). Predator odor increases avoidance and glutamatergic synaptic transmission in the prelimbic cortex via corticotropin-releasing factor receptor 1 signaling. Neuropsychopharmacology, 44(4), 766–775. doi:10.1038/s41386-018-0279-2

Jaramillo, A. A., Randall, P. A., Frisbee, S., & Besheer, J. (2016). Modulation of sensitivity to alcohol by cortical and thalamic brain regions. Eur J Neurosci, 44(8), 2569–2580. doi:10.1111/ejn.13374

Jayachandran, M., Linley, S. B., Schlecht, M., Mahler, S. V., Vertes, R. P., & Allen, T. A. (2019). Prefrontal Pathways Provide Top-Down Control of Memory for Sequences of Events. Cell Rep, 28(3), 640–654 e646. doi:10.1016/j.celrep.2019.06.053

Kafetzopoulos, V., Kokras, N., Sotiropoulos, I., Oliveira, J. F., Leite-Almeida, H., Vasalou, A., Sardinha, V. M., Papadopoulou-Daifoti, Z., Almeida, O. F. X., Antoniou, K., Sousa, N., & Dalla, C. (2018). The nucleus reuniens: a key node in the neurocircuitry of stress and depression. Mol Psychiatry, 23(3), 579–586. doi:10.1038/mp.2017.55

Kessler, R. C., Berglund, P., Demler, O., Jin, R., Merikangas, K. R., & Walters, E. E. (2005). Lifetime prevalence and age-of-onset distributions of DSM-IV disorders in the National Comorbidity Survey Replication. Arch Gen Psychiatry, 62(6), 593–602. doi:10.1001/archpsyc.62.6.593

Kessler, R. C., Sonnega, A., Bromet, E., Hughes, M., & Nelson, C. B. (1995). Posttraumatic stress disorder in the National Comorbidity Survey. Arch Gen Psychiatry, 52(12), 1048–1060. doi:10.1001/archpsyc.1995.03950240066012

Kilpatrick, D. G., Resnick, H. S., Milanak, M. E., Miller, M. W., Keyes, K. M., & Friedman, M. J. (2013). National estimates of exposure to traumatic events and PTSD prevalence using DSM-IV and DSM-5 criteria. J Trauma Stress, 26(5), 537–547. doi:10.1002/jts.21848

Lin, Y. J., Chiou, R. J., & Chang, C. H. (2020). The Reuniens and Rhomboid Nuclei Are Required for Acquisition of Pavlovian Trace Fear Conditioning in Rats. eNeuro, 7(3). doi:10.1523/ENEURO.0106-20.2020

Maisson, D. J., Gemzik, Z. M., & Griffin, A. L. (2018). Optogenetic suppression of the nucleus reuniens selectively impairs encoding during spatial working memory. Neurobiol Learn Mem, 155, 78–85. doi:10.1016/j.nlm.2018.06.010

Mei, H., Logothetis, N. K., & Eschenko, O. (2018). The activity of thalamic nucleus reuniens is critical for memory retrieval, but not essential for the early phase of "off-line" consolidation. Learn Mem, 25(3), 129–137. doi:10.1101/lm.047134.117

Neal, S., Kent, M., Bardi, M., & Lambert, K. G. (2018). Enriched Environment Exposure Enhances Social Interactions and Oxytocin Responsiveness in Male Long-Evans Rats. Front Behav Neurosci, 12, 198. doi:10.3389/fnbeh.2018.00198

Ornelas, L. C., Tyler, R. E., Irukulapati, P., Paladugu, S., & Besheer, J. (2020). Increased alcohol self-administration following exposure to the predator odor TMT in active coping female rats. Behav Brain Res, 402, 113068. doi:10.1016/j.bbr.2020.113068

Ramanathan, K. R., Jin, J., Giustino, T. F., Payne, M. R., & Maren, S. (2018). Prefrontal projections to the thalamic nucleus reuniens mediate fear extinction. Nat Commun, 9(1), 4527. doi:10.1038/s41467-018-06970-z

Ramanathan, K. R., & Maren, S. (2019). Nucleus reuniens mediates the extinction of contextual fear conditioning. Behav Brain Res, 374, 112114. doi:10.1016/j.bbr.2019.112114

Ramanathan, K. R., Ressler, R. L., Jin, J., & Maren, S. (2018). Nucleus Reuniens Is Required for Encoding and Retrieving Precise, Hippocampal-Dependent Contextual Fear Memories in Rats. J Neurosci, 38(46), 9925–9933. doi:10.1523/JNEUROSCI.1429-18.2018

Randall, P. A., Lovelock, D. F., VanVoorhies, K., Agan, V. E., Kash, T. L., & Besheer, J. (2021). Low-dose alcohol: Interoceptive and molecular effects and the role of dentate gyrus in rats. Addict Biol, 26(3), e12965. doi:10.1111/adb.12965

Riittinen, M. L., Lindroos, F., Kimanen, A., Pieninkeroinen, E., Pieninkeroinen, I., Sippola, J., Veilahti, J., Bergstrom, M., & Johansson, G. (1986). Impoverished rearing conditions increase stress-induced irritability in mice. Dev Psychobiol, 19(2), 105–111. doi:10.1002/dev.420190203

Rosen, J. B., Asok, A., & Chakraborty, T. (2015). The smell of fear: innate threat of 2,5-dihydro-2,4,5-trimethylthiazoline, a single molecule component of a predator odor. Front Neurosci, 9, 292. doi:10.3389/fnins.2015.00292

Schwendt, M., Shallcross, J., Hadad, N. A., Namba, M. D., Hiller, H., Wu, L., Krause, E. G., & Knackstedt, L. A. (2018). A novel rat model of comorbid PTSD and addiction reveals intersections between stress susceptibility and enhanced cocaine seeking with a role for mGlu5 receptors. Transl Psychiatry, 8(1), 209. doi:10.1038/s41398-018-0265-9

Shallcross, J., Hamor, P., Bechard, A. R., Romano, M., Knackstedt, L., & Schwendt, M. (2019). The Divergent Effects of CDPPB and Cannabidiol on Fear Extinction and Anxiety in a Predator Scent Stress Model of PTSD in Rats. Front Behav Neurosci, 13, 91. doi:10.3389/fnbeh.2019.00091

Silva, B. A., Burns, A. M., & Graff, J. (2019). A cFos activation map of remote fear memory attenuation. Psychopharmacology (Berl), 236(1), 369–381. doi:10.1007/s00213-018-5000-y

Tyler, R. E., Weinberg, B. Z. S., Lovelock, D. F., Ornelas, L. C., & Besheer, J. (2020). Exposure to the predator odor TMT induces early and late differential gene expression related to stress and excitatory synaptic function throughout the brain in male rats. Genes Brain Behav, 19(8), e12684. doi:10.1111/gbb.12684

Weera, M. M., Schreiber, A. L., Avegno, E. M., & Gilpin, N. W. (2020). The role of central amygdala corticotropin-releasing factor in predator odor stress-induced avoidance behavior and escalated alcohol drinking in rats. Neuropharmacology, 166, 107979. doi:10.1016/j.neuropharm.2020.107979

Xu, W., & Sudhof, T. C. (2013). A neural circuit for memory specificity and generalization. Science, 339(6125), 1290–1295. doi:10.1126/science.1229534

Yehuda, R. (2004). Risk and resilience in posttraumatic stress disorder. J Clin Psychiatry, 65 Suppl 1, 29– 36.

